# Tad pili play a dynamic role in *Caulobacter crescentus* surface colonization

**DOI:** 10.1101/526160

**Authors:** Matteo Sangermani, Isabelle Hug, Nora Sauter, Thomas Pfohl, Urs Jenal

**Affiliations:** Biozentrum, University of Basel, Klingelbergstrasse 50/70, 4056 Basel, Switzerland.; Department of Chemistry, University of Basel, Klingelbergstrasse 80, 4056 Basel, Switzerland.; Swiss Nanoscience Institute, 4056 Basel, Switzerland.; Institute of Physics, University of Freiburg, Hermann-Herder-Str 3, 79104 Freiburg, Germany.

**Keywords:** Tad pili, Type IV pili, *Caulobacter crescentus*, surface attachment, microfluidics, c-di-GMP, mechanosensation

## Abstract

Bacterial surface attachment is mediated by rotary flagella and filamentous appendages called pili. Here, we describe the role of Tad pili during surface colonization of *Caulobacter crescentus*. Using an optical trap and microfluidic controlled flow conditions as a mimic of natural environments, we demonstrate that Tad pili undergo repeated cycles of extension and retraction. Within seconds after establishing surface contact, pili reorient cells into an upright position promoting walking-like movements against the medium flow. Pili-mediated positioning of the flagellated pole close to the surface facilitates motor-mediated mechanical sensing and promotes anchoring of the holdfast, an adhesive substance that affords long-term attachment. We present evidence that the second messenger c-di-GMP regulates pili dynamics during surface encounter in distinct ways, promoting increased activity at intermediate levels and retraction of pili at peak concentrations. We propose a model, in which flagellum and Tad pili functionally interact and together impose a ratchet-like mechanism that progressively drives *C. crescentus* cells towards permanent surface attachment.

## INTRODUCTION

Bacteria have evolved effective mechanisms to colonize abiotic and biotic surfaces in order to scavenge nutrients, attack host tissue or assemble into refractory communities called biofilms. A pivotal role in this process is played by adhesive pili, also called fimbriae, protein-based filaments exposed on the surface of bacteria that have adopted different functions including adherence, motility, electron transfer, acquisition of DNA and protein secretion (Giltner *et al*, 2012; Hospenthal *et al*, 2017). Accordingly, pili are crucial virulence factors during infection processes (Craig *et al*, 2004). They mediate direct contact between pathogens and specific host tissues and promote pathogen spreading and cellular invasion (Merz *et al*, 2000; Skerker & Berg, 2001; Chang *et al*, 2017a, 2017b; Gold *et al*, 2015). The highly corrugated surface of the extended filaments mediates attachment to hydrophobic and hydrophilic surfaces through reversible, nonspecific interactions (Lu *et al*, 2015; Kachlany *et al*, 2000). The most sophisticated class of these filaments are Type IV pili. The best studied subgroup, Type IVa, are dynamic machineries that undergo cycles of extension and retraction through the rapid assembly and disassembly of pilin subunits at the proximal end of the structure (Chang *et al*, 2017a; McCallum *et al*, 2017). Extension and retraction are powered by specific cytoplasmic ATPases, which generate rotational movements of the assembly platform in the inner membrane to incorporate pilin subunits into or extract them from the helical filaments (Merz *et al*, 2000; Pelicic, 2008). Through the coordinated extension and retraction of multiple polar pili, single cells are able to move on surfaces and explore their environments (Giltner *et al*, 2012; Berry & Pelicic, 2015; Schilling *et al*, 2010). Two types of pili-mediated movements have been described (Maier & Wong, 2015). Crawling movements of horizontally positioned cells, called twitching, can cover large distances with high directional persistence (Jin *et al*, 2011; Sun *et al*, 2000; Skotnicka *et al*, 2016). In contrast, walking movements of orthogonal upright cells facilitates rapid exploration of smaller areas (Conrad *et al*, 2011).

Type IV pili are widespread in bacteria and archaea (Pelicic, 2008; Berry & Pelicic, 2015). Distinctive features divide these structures into two classes, called Type IVa and Type IVb. Type IVa represents a uniform class that is found in important human pathogens like *Pseudomonas aeruginosa*, *Vibrio cholerae* or *Neisseria* spp. and in environmental bacteria like *Myxococcus xanthus*, *Shewanella putrefaciens* or *Bdellovibrio bacteriovorus*. The Type IVb subclass is less homogenous and best-characterized for enteropathogenic *Escherichia coli* or *V. cholerae* (Tomich *et al*, 2007). A subclass of Type IVb pili are the tight adherence (Tad) or Flp (fimbrial low-molecular-weight protein) pili (Figure 1A) (Skerker & Shapiro, 2000; Kachlany *et al*, 2001; Mignolet *et al*, 2018). Sometimes classified as a separate Type IVc group, Tad pilin subunits are shorter as compared to pilin of other Type IV systems, but show similar hydrophobic intermolecular interactions providing the main force holding the fibers together (Tomich *et al*, 2007; Kachlany *et al*, 2001; Schilling *et al*, 2010). Tad pili promote surface colonization, cell-to-cell aggregation, biofilm cohesion and promote virulence of different bacterial pathogens (Tomich *et al*, 2007; Kachlany *et al*, 2001; Schilling *et al*, 2010; Wang *et al*, 2014; Nykyri *et al*, 2013; Bernier *et al*, 2017). In contrast to Type IVa and other Type IVb pili systems, Tad clusters lack a dedicated retraction ATPase and most Tad pili do not seem to be dynamic (Tomich *et al*, 2007). However, recent studies indicated that toxin-coregulated pili (TCP) of *V. cholerae* and Tad pili of *C. crescentus* are retractable (Ng *et al*, 2016; Ellison *et al*, 2017). For TCP it was proposed that incorporation of minor pilin subunits, TcpB, into a growing pilus can block assembly and trigger a spontaneous disassembly and retraction event.

**FIGURE 1.**
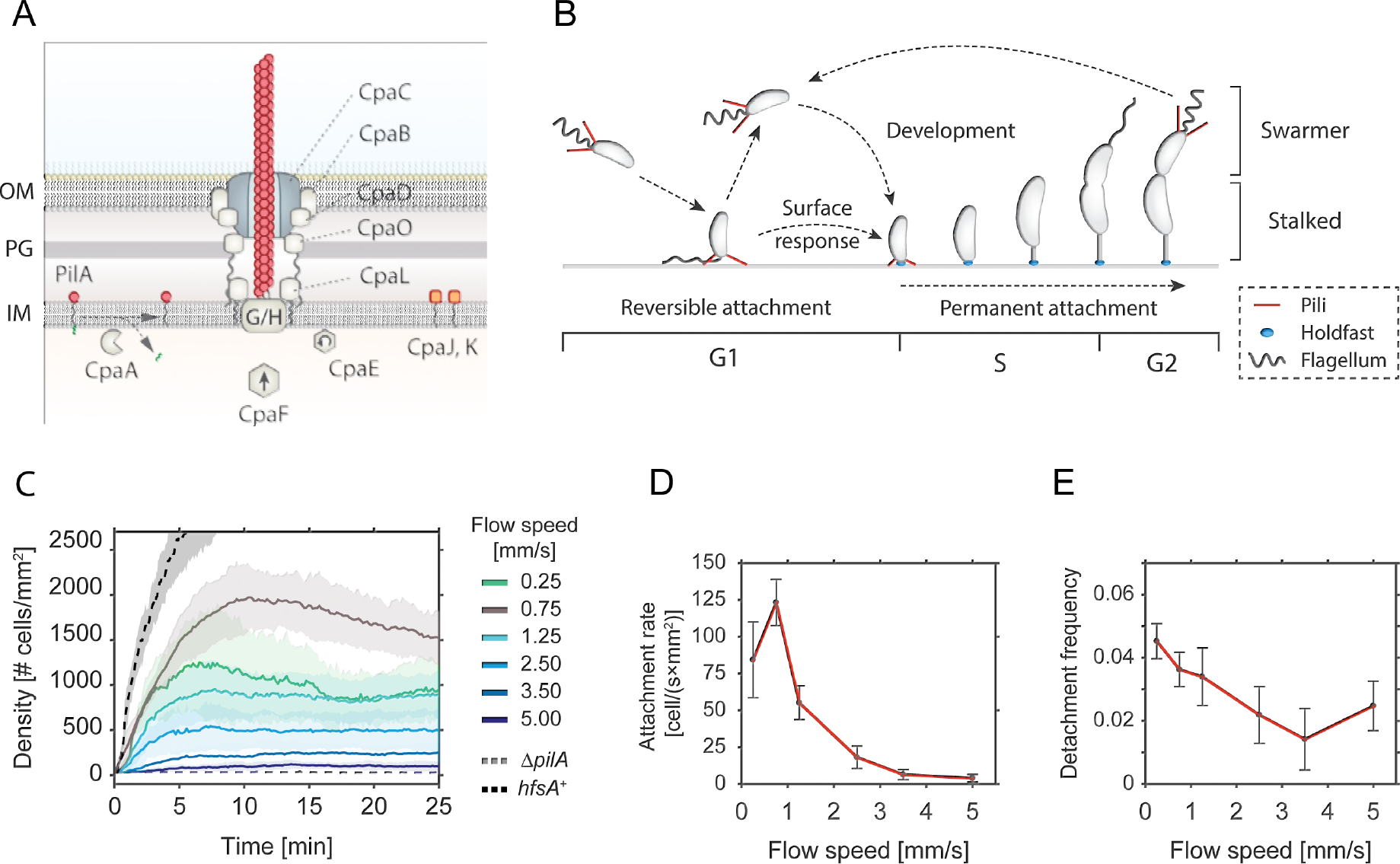
Pili-mediated surface attachment of *C. crescentus* cells in flow conditions. **A -** Schematic model of the Tad pili machinery in *C. crescentus*. The function of individual components is drawn according to comparative analysis of the cpa locus with other Tad pili (Tomich *et al*, 2007; Mignolet *et al*, 2018). PilA is the major pilin subunit that matures upon removal of the signal peptide by the prepilin peptidase CpaA and is then assembled into a filament by the inner membrane platform that is located at the base of the filament (CpaG and CpaH). The CpaF ATPase is the functional motor protein, while the CpaE ATPase is required for polar localization of the pilus machinery (Tomich *et al*, 2007; Mignolet *et al*, 2018). Minor pilin subunits (CpaJ,K) and envelope spanning components of the pili machinery are indicated. **B -** Schematic of the *C. crescentus* cell cycle. SW cells are born with assembled pili (red) and flagellum (grey). Upon surface encounter of the SW cell, pili promote temporary attachment and position the flagellated cell pole close to the surface. This triggers the secretion of an adhesive exopolysaccharide, the holdfast (blue), and results in permanent attachment of the cell. Attached cells differentiate into ST cells, initiate the division cycle that generates another motile SW cell. **C -** Flow velocity of the fluid influences pili-mediate surface attachment. The chart shows numbers of *C. crescentus* cells that lack an adhesive holdfast (*hfsA*^−^) adhering to the flow channel surface at different flow velocities. Experiments with a wild-type strain able to produce holdfast (*hfsA*^+^) and with a mutant lacking pili (Δ*pilA*) are shown as controls at a flow speed of 0.75 mm/s. Opaque areas represent standard deviations (*n*>3). **D -** Average number of newly attached SW cells per mm^2^ and second at different flow velocities. The values are averages from the microfluidic attachment assay in Figure 1C scored between 10 and 25 minutes. Error Bars represent standard deviations (*n*>3). **E -** Frequency of surface detachment at different flow velocities. Detachment frequencies represent the ratio between the number of cells leaving the surface over the total number attached. The values are averages from the microfluidic attachment assay in Figure 1C scored between 10 and 25 minutes. Error Bars represent standard deviations (*n*>3).

In *C. crescentus* polar Tad pili facilitate the attachment of motile planktonic cells to surfaces (Hug *et al*, 2017; Ellison *et al*, 2017; Skerker & Shapiro, 2000). During *C. crescentus* division, a polarized sessile stalked cell (ST) produces a motile offspring, the swarmer cell (SW), which is equipped with a single flagellum and multiple pili (Figure 1B) (Viollier *et al*, 2002; Skerker & Shapiro, 2000). The new-born SW cell remains in a motile, non-replicating state for a defined period called G1. After this period, chromosome replication resumes coincident with cell differentiation, during which flagellum and pili are replaced by an adhesive exopolysaccharide, the holdfast, and the stalk. While the developmental program defines an extended time window of motility, SW cells that are challenged with surface are able to transit to the sessile state within seconds (Hug *et al*, 2017; Li *et al*, 2012; Persat *et al*, 2014). This process is executed by a surface recognition program that involves sensing of mechanical cues and triggering of a burst of c-di-GMP, a second messenger that controls the motile-sessile transition in a wide range of bacteria (Abel *et al*, 2013; Conner *et al*, 2017; Luo *et al*, 2015). In turn, c-di-GMP allosterically activates a pre-assembled holdfast synthesis machinery to irreversibly anchor cells on the surface (Hug *et al*, 2017; Sprecher *et al*, 2017). However, while the initial surface contact and adherence is indisputably pili-mediated, different views have been put forward for the role of polar Tad pili in *C. crescentus* surface sensing. In one model, pili activity positions the flagellar pole in close contact with the surface to allow mechanosensation by the membrane-integral rotary motor triggering a spike of c-di-GMP through the activation of a motor-associated diguanylate cyclase (Hug *et al*, 2017). A second model proposed that surface-bound Tad pili themselves serve as surface sensors, mediating an internal upshift of c-di-GMP after experiencing resistance on retracting (Ellison *et al*, 2017). In line with the first model, recent studies demonstrated that the flagellar motor, but not Type IV pili are required for surface sensing in *P. aeruginosa* (Laventie *et al*, 2018) and that a motor-coupled diguanylate increases levels of c-di-GMP in this organism (Baker *et al*, 2019). Moreover, studies in *V. cholerae* and *P. aeruginosa* had shown that c-di-GMP is positioned upstream of Type IV pili and that this second messenger regulates pili assembly and activity (Guzzo *et al*, 2009; Jain *et al*, 2017; Jones *et al*, 2015; Laventie *et al*, 2018).

To more scrutinize the role of Tad pili in *C. crescentus* surface recognition and surface colonization, we carefully analysed their dynamic behaviour and regulation. We used microfluidics to perform experiments under controlled flow conditions. We demonstrate that under steady medium flow, Tad-mediated cell attachment is transient, offering motile bacteria a short window of opportunity during which they can trigger holdfast biogenesis. Tad pili are highly dynamic already before the motile SW separates from its ST mother, explaining the observed ultra-rapid surface recognition of new-born SW cells (Hug *et al*, 2017). We show that Tad pili can go through multiple rounds of extension and retraction mediating walking-like motility on surfaces. Finally, we present data indicating that the flagellar motor and Tad pili functionally interact and that an increase of c-di-GMP concentration results in pili retraction. Together, these data propose a model, in which Tad pili structures are highly dynamic and represent an integrated part of a complex mechanism that senses mechanical stimuli upon *C. crescentus* surface encounter to promote and accelerate surface anchoring.

## RESULTS

### Tad pili mediate transient surface attachment under media flow

Surface adhesion of *C. crescentus* via its polar pili is transient and weaker as compared to the strong and long-lasting attachment via the adhesive holdfast (Berne *et al*, 2013). To investigate the overall contribution of pili to surface attachment without interference of the holdfast, we analysed the behaviour of a *C. crescentus* holdfast mutant (NA1000 *hfsA*^−^) (Marks *et al*, 2010) and scored attachment efficiency in simple microfluidic channels with a single inlet supplying a culture with a constant flow of medium. Microscopy time lapse images were recorded to determine the rate of surface attachment. Importantly, mutants lacking pili were unable to adhere to the glass surface in such an assay (Figure 1C).

The number of attached cells scaled with the flow velocity of the medium (Figure 1C). We observed a plateau in the density of attached cells per unit surface area, arguing that attachment is transient and that, under these conditions, attachment and detachment of bacteria reached an equilibrium (Supplemental Movie 1). A plateau was generally reached 5-10 min after the start of the experiment. Colonization density at equilibrium showed a strong dependency on the flow velocity. The highest density of attached cells was observed with a fluid flow velocity in the microchannel of 0.75 mm/s (maximal velocity in the middle of the channel). This velocity was chosen as standard for further experiments. At lower flow speed we measured a lower plateau value of colonization density (Figure 1C) due to a decreased attachment rate of cells, while the detachment frequency was unchanged as compared to the optimal flow velocity of 0.75 mm/s (Figure 1D and 1E). Similarly, with increasing flow velocities, colonization densities also decreased with attachment being completely abolished above 5 mm/s cell (Figure 1C). The decreased plateau levels were primarily due to a 2 to 10-fold decrease in attachment rates, while the detachment frequencies were 2 to 3-fold lower, meaning attached cells were less likely to leave (Figure 1D and 1E). As a consequence, the average residence time of piliated SW cells on glass surface was increased at higher flow (Supplemental Figure 1A). E.g. at a flow of 0.75 mm/s 30% of the cells were retained on the surface for more than two minutes, while this fraction increased to 50-60% at higher flow rates.

Surface bound cells subjected to fluid flow experience a drag proportional to the flow velocities (Supplemental Figure 1C). The observed increase in residence time at higher flow velocities may result from conformational changes in surface attached pili fibers that strengthen the interaction with the surface at higher drag forces (Kolappan *et al*, 2016; Biais *et al*, 2010). Together, these observations demonstrated that pili mediate transient surface attachment of *C. crescentus* SW cells under media flow. Importantly, pili negotiate a time window of a few minutes for tactile sensing and holdfast production, to undergo the transition from temporary to permanent attachment (Hug *et al*, 2017). In agreement with this view, cells harboring an intact holdfast machinery (*hfsA*^+^) rapidly attach to and permanently colonize glass surfaces (Figure 1C).

### Tad pili are active in the predivisional cell before division

Surface attached *C. crescentus* SW cells were most often found standing upright perpendicular to the surface, irrespective of their ability to synthesize an adhesive holdfast (Supplemental Movies 1, 2) (Hug *et al*, 2017). This is an unfavorable position considering the constant drag force from the media flow. Also, after landing, cells occasionally moved a few microns against the medium flow in an upright position. Since only an active force could move cells against or position them perpendicular to the flow, we speculated that surface bound Tad pili are able to retract under these conditions (Conrad *et al*, 2011). To investigate pili dynamics we first analysed strains that are capable of secreting an adhesive holdfast in flow channels mimicking conditions that *C. crescentus* encounters in its natural environment (Persat *et al*, 2014). Under such conditions, offspring of attached dividing mothers are exposed to surface before division as a consequence of medium flow over the crescentoid dividing cells (Figure 2) (Persat *et al*, 2014).

**FIGURE 2.**
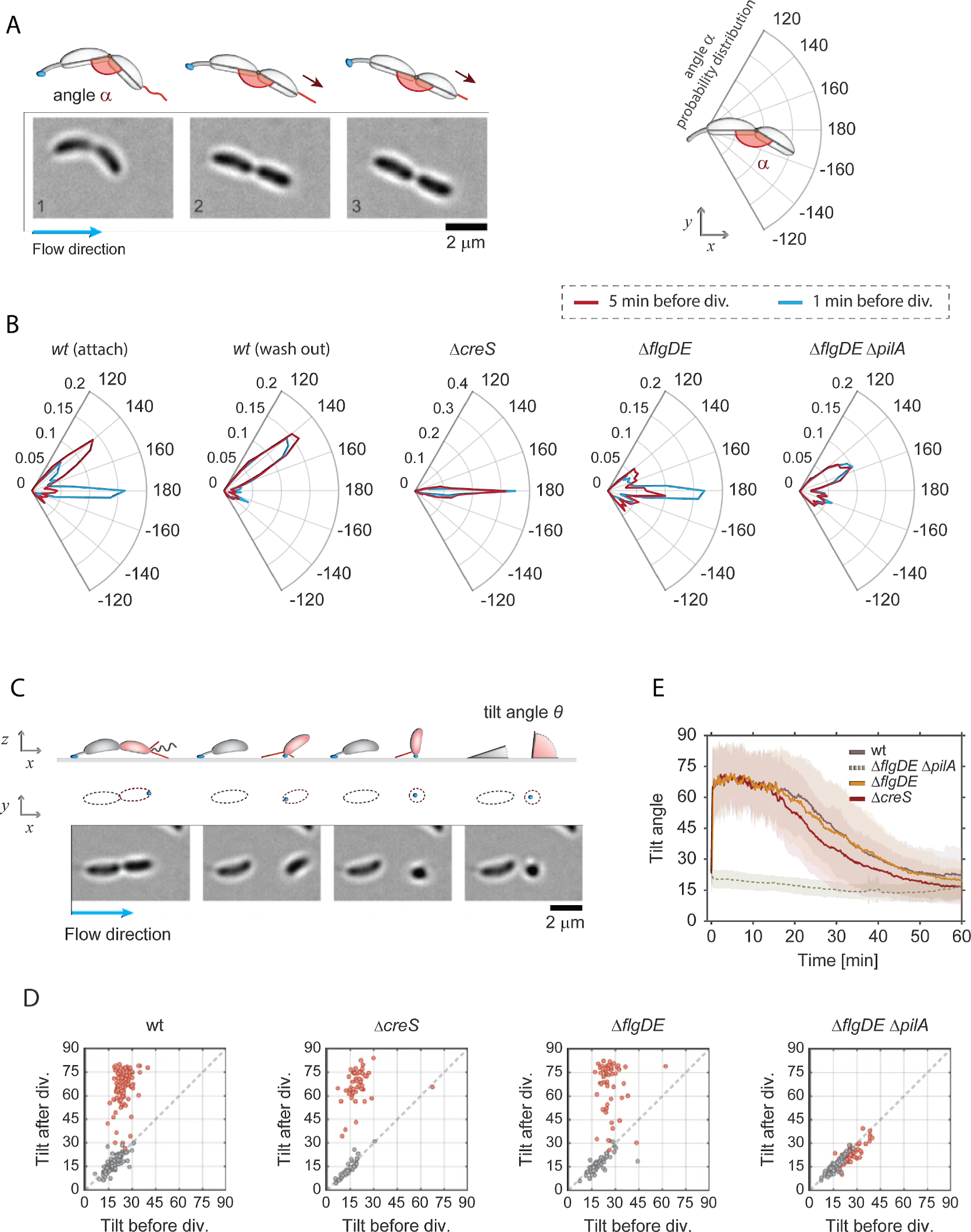
Pili are active before cell separation and afterwards position surface-bound cells upright. **A -** Image sequence of a crescentoid predivisional cell that is attached via its holdfast at one pole and is stretching due to the activity of pili located at the opposite pole. A schematic representation of the cell with its holdfast (blue) and pili (red) is shown above the micrographs, illustrating how the angle *α* was determined in each experiment. The direction of medium flow is indicated. **B -** Angle distribution along the main axis of predivisional cells. A schematic of a predivisional cell with the angle between ST and SW progeny is shown on the top right of the polar charts. Each plot shows the distribution of angle *α* in a different strain recorded five minutes (dark red) and one minute (blue) before cell division. The *C. crescentus* wild type strain has a peak at about 150° resulting from the crescentoid shape of predivisional cells at rest. A second peak is observed at 180° resulting from pili retraction and cell stretching into a straight line. Controls lacking pili (Δ*pilA*), crescentin (Δ*creS*) or the flagellum (Δ*flgDE*) are shown. Stretching of dividing cells that retained the SW offspring on the surface after separation (“attach”) and cell that failed to attach (“wash out”) are indicated. Replicas: wt (attach) = 130, wt (wash out) = 119, Δ*creS* = 56, Δ*flgDE* = 57, Δ*flgDE* Δ*pilA* = 75. **C -** Time-lapse images of *C. crescentus* cell division under flow (bottom) with a schematic view (top). A ST mother cell attached to the surface via its holdfast (blue) produces a SW offspring (red). A schematic representation of the cell outline in the xy plane is shown as identified by the analysis program. Polar pili are highlighted (red). Newborn SW cells but not ST mothers are able to move into a vertical position after separation. The tilt angles ϑ (angle between the main cell body axis and the glass surface) were calculated from the cell contour shape (*xy* plane) with cells lying parallel to the glass surface and being in an upright position scoring ϑ=0° and ϑ=90°, respectively. The direction of medium flow is indicated. **D -** Pili are required for newborn SW cells to move into an upright position. Scatter plots comparing the average angle ϑ of the same cells recorded 5 minutes before and 5 minutes after cell division. The results shown were obtained with the strains indicated. SW cells are in red and data for ST cells are in grey. **E -** Dynamics of pili-mediated standing up of SW cells. The change of the angle ϑ after birth (time point 0) of SW cells is shown over time for *C. crescentus* wild type, Δ*creS* and Δ*flgDE* Δ*pilA* mutants. Please note that the Δ*pilA* control strain also contained a Δ*flgDE* deletion, as cells lacking the external parts of the flagellum show hypersensitive surface response (Hug *et al*, 2017) and thus allowed to trap enough pili mutant cells on surfaces for this analysis. Solid lines represent average values; opaque areas show standard deviations. Replicas in D and E: wt = 98; Δ*flgDE* = 57; Δ*creS* = 56; Δ*flgDE* Δ*pilA* = 75.

To investigate if pili are active already before cells divide, movements of surface attached dividing cells were carefully analysed. We observed that the piliated pole of late predivisional cells was pulled away from the ST pole, stretching the typical crescentoid shape into a straight line (Figure 2A, Supplemental Movie 2). To quantify this behaviour, we determined the angle (*α*) between the two cell bodies five minutes and one minute prior to cell separation. *C*-shaped *C. crescentus* cells with pili showed discrete peaks of angle *α* at 180° and at 150° (Figure 2B). In comparison, a Δ*creS* mutant that lacks the characteristic crescentoid cell curvature (Ausmees *et al*, 2003) showed only a single peak at 180°. This indicated that the peak at 150° represents cells with their natural, unstrained crescentin-mediated curvature in the late predivisional stage. A mutant lacking pili did not show a peak at 180°, but retained peaks at 150°, arguing that cell stretching before division is mediated by the action of polar pili. Consistent with this, cells lacking the polar flagellum retained their stretching ability (Figure 2B). Moreover, cell stretching was more prominent at very late stages of division, with the peak at 180° increasing at the expense of the peak at 150° (Figure 2B). Finally, we observed a striking difference between the behaviour of predivisional cells that gave birth to offspring able to attach after separation (Figure 2B, attach) and predivisional cells producing SW cells that were washed out after cell division (Figure 2B, wash out). While the former showed a prominent peak at 180°, this peak was missing in dividing cells destined to produce offspring unable to attach. Instead, the latter showed a distribution of angle *α* resembling the pilus-deficient strain (Figure 2B). The correlation of pili-mediated cell stretching and successful surface attachment of SW offspring suggested that pili play an active role in surface attachment. This is consistent with the observation that SW offspring freely rotating before detachment from their mothers, generally failed to remain attached to the surface (Hug *et al*, 2017). Of note, the orientation of the concavity of attached predivisional cells showed a strong bias to one side with respect to the flow direction (Figure 2B, Supplemental Figures 2A-C). This suggested that the *C*-shape of *C. crescentus* cells has a small helical twist (Supplemental Figure 2). While the concavity distribution is expected to be random for cells with a straight *C*-shape, we found that cells were more likely to orient to the left with respect to the flow direction, irrespective of their position within the channel or the nature of the surface.

Together, these experiments indicated that pili are assembled and active at the flagellated pole before cell division takes place and that pili activity increases as dividing cells approach the separation stage. Moreover, pili activity is instrumental for surface attachment. We had proposed earlier that pili retraction at this stage of cell division is critical to position the flagellar mechanosensor in close proximity to the surface in order to successfully initiate biogenesis of the adhesive holdfast and therefore preventing cells from being washed out (Hug *et al*, 2017).

### Dynamic Tad pili pull attached swarmer cells into an upright position

We observed that within a few seconds after cell division, new-born SW cells were standing up against the medium flow (Supplemental Movie 2). To quantify these movements, we measured the 2D projections of individual cells in the *xz* plane and used this information to infer the cells’ 3D orientation and tilt angles (ϑ) over time (Figure 2C). New-born SW cells were unable to change their position as long as they were physically connected to their ST mothers. However, upon separation, SW cells rapidly changed their tilt, moving into an upright position of about 60-75° degrees. In contrast, ST cells retained their low ϑ value after cell division (Figure 2D and 2E). Because mutants lacking pili (Δ*pilA*) showed very low attachment, we analysed cell movements of a strain lacking both the major pilin subunit as well as the distal parts of the polar flagellum (Δ*flgDE*) as a control. We had shown earlier that *C. crescentus* cells lacking the outer parts of the polar flagellum (Δ*flgDE*) show a hypersensitive surface response that partially alleviates the strict requirement for pili (Hug *et al*, 2017). In agreement with this, SW cells of the Δ*flgDE* Δ*pilA* mutant sporadically remained surface attached after separation from their mothers. Importantly, all cells invariably retained the same low Ñ angle value (Figure 2D and Supplemental Movie 3). In contrast, a strain lacking only the flagellar structures (Δ*flgDE*) showed wild type-like standing-up behaviour (Figure 2D and 2E). Deleting the crescentin cytoskeleton (Δ*creS*) did not influence the ability of attached cells to stand up, arguing that cell curvature does not influence dynamic cell movements after division. Together, this indicated that force-generating pili are required for newborn SW cells to immediately move into a vertical, upright position. Despite the relatively strong medium flow, daughter cells were able to keep their upright position for about 10-15 minutes before the angle ϑ gradually decreased. This coincides well with the timing of the swarmer-to-stalked cell differentiation when pili disappear and stalk biogenesis is initiated (Figure 1B).

### Tad pili mediate walking movements against the media flow

Generally, pili-mediated cell movements into an upright position or predivisional cell stretching were observed only once while repeated cycles of extension and retraction were observed occasionally. Moreover, in holdfast deficient cells we occasionally observed cells moving for very short distances (1-3 μm) after landing on surface. We reasoned that prevailing activity of pili might be masked in attached SW cells by the rapid synthesis of the holdfast adhesin, which immobilizes cells in an upright position (Hug *et al*, 2017). To address if *C. crescentus* pili are capable of undergoing multiple cycles of extension and retraction, we made use of a *hfsK* mutant (Sprecher *et al*, 2017). *HfsK* is a c-di-GMP effector protein, the activity of which increases holdfast cohesion. Mutants lacking this protein secrete holdfast material that is strong enough to glue cells to the surface in microfluidic devices with media flow, but fails to firmly anchor cells at the place of initial attachment (Sprecher *et al*, 2017). To tune the amount of holdfast generated, a Δ*hfsK* strain was engineered that allowed modulating intracellular levels of c-di-GMP (*rcdG*^0^::*P*_*lac*_-*dgcZ*), the primary allosteric regulator of the holdfast secretion machinery, via the controlled expression of the exogenous diguanylate cyclase gene *dgcZ* (Hug *et al*, 2017). In flow chambers, cells of this strain were dragged across the surface by the media flow leaving trails of stained holdfast material behind (Sprecher *et al*, 2017). Intriguingly, we observed that new-born SW cells, after standing up, were able to move upstream against the medium flow (Figure 3A, B and Supplemental Movie 4). These results indicated that pili can retain their dynamic behaviour to dislodge cells that are weakly attached to the surface. Tracking the trajectories of individual cells identified the time period, step size, and speed of pili-driven movements (Figure 3B). On average, moving cells reached a speed of about 300 nm/sec covering distances of up to 500 nm in individual steps (Figure 3C and Supplemental Figure 3A). Overall, cells covered distances of several micrometers in repetitive small steps with a frequency of about 2-3 steps per minute (Figure 3C). While the step frequency remained high for the first 10-15 minutes after division, it gradually decreased over time and discontinued about 20-30 min after daughters had separated from their mothers (Figure 3C, Supplemental Movie 4). During their movements, SW cells, although unable to remain standing for longer periods of time, repeatedly moved back into an upright position (Figure 3D). This behaviour was particularly pronounced at the conclusion of each step event, arguing that full retraction of pili forces cells into an upright position (Figure 3D and Supplemental Figure 3B). Together, these results indicated that *C. crescentus* pili remain highly active over longer periods of time and that, under these conditions, they engage in repetitive cycles of extension and retraction. This is highly reminiscent of twitching or walking movements observed for other Type IV pili-positive bacteria (Conrad *et al*, 2011).

**FIGURE 3.**
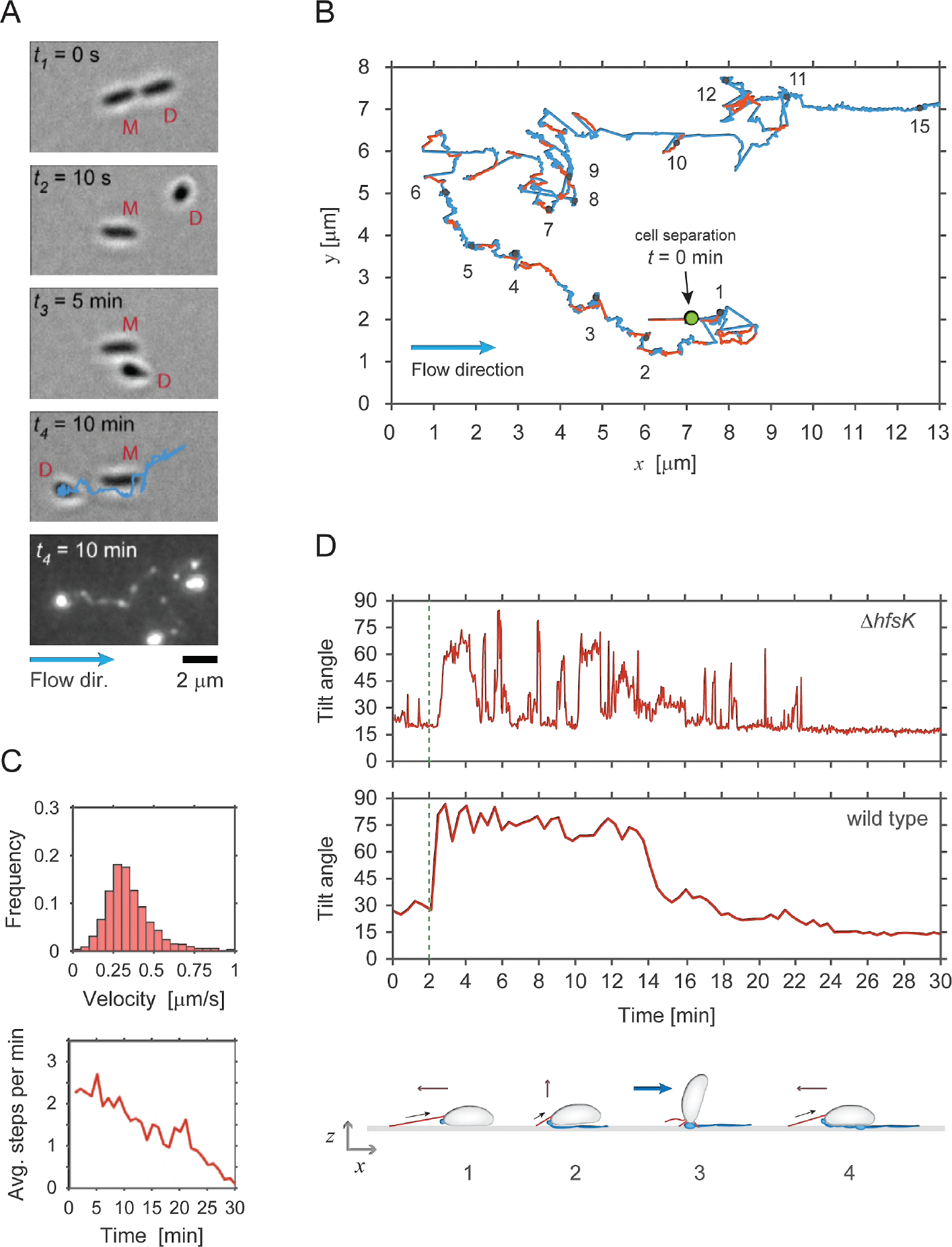
Dynamic pili assist walking-like movements against the medium flow. **A -** Example of a newborn SW cell of a Δ*hfsK* mutant moving against the medium flow. The secretion of holdfast adhesin is monitored microscopically by employing fluorescently labelled wheat germ agglutinin (bottom image). The time after cell division is indicated and mother (M) and daughter (D) cells are individually labelled. Time points *t*_1_ and *t*_2_ show the SW cell immediately before and after separation from its mother. Time points *t*_3_ and *t*_4_ show how the SW cell moves against the medium flow (blue arrow) past its mother. The blue track, which indicates the trajectory of the cell recorded during its 10 min walk, perfectly matches trails of holdfast material left behind. **B -** Representative trajectory (blue line) of a SW cell moving on the surface of a microfluidic channel. The trajectory is reconstructed from time-lapse images recorded for a single cell of the Δ*hfsK* mutant. Step events were identified as fast movements against the flow and are highlighted in red. Black dots in the track indicate the time (minutes) after cell separation. **C -** Pili-mediated walking speeds (Upper chart) and average number of step events per minute (lower chart) recorded for SW cells of the Δ*hfsK* mutant at the time points indicated after cell division (time zero) (*n* = 56). **D -** SW cells are repeatedly standing up during movements against medium flow. The tilt angle ϑ was recorded over time in representative examples of walking SW cells of a Δ*hfsK* mutant (upper panel) and wild type (lower panel), respectively. A schematic of walking movements of the Δ*hfsK* mutant is shown below the charts. Retraction of an extended pilus pulls a horizontally positioned cell forward (1). Upon full retraction of the pilus, the cell body is pulled into an upright position (2), against the drag force of the medium flow (3). Upon completion of pilus retraction, the cell is pushed back onto the surface by flow (4) followed by the next motility step catalysed by an extended pilus.

### Determining the force and energy of pili retraction

To measure the forces generated by pili retraction we made use of optical tweezers. Because *C. crescentus* cells are highly susceptible to phototoxicity elicited by the strong light source used in optical traps, it was not practical to directly manipulate cells with this method. To avoid this problem, we exploited the ability of *C. crescentus* predivisional cells to permanently adhere to surfaces via their adhesive holdfast. By coupling predivisional cells to polystyrene beads and immobilizing the beads in the optical trap, phototoxicity was eliminated, as cells could now be kept in the optical trap for at least 1 hour without losing their ability to grow and divide (Figure 4A). By moving the beads carrying predivisional cells towards the glass surface, pili were able to attach and, upon retraction, displace the beads from the trap (Figure 4A and Supplemental Movie 5).

**FIGURE 4.**
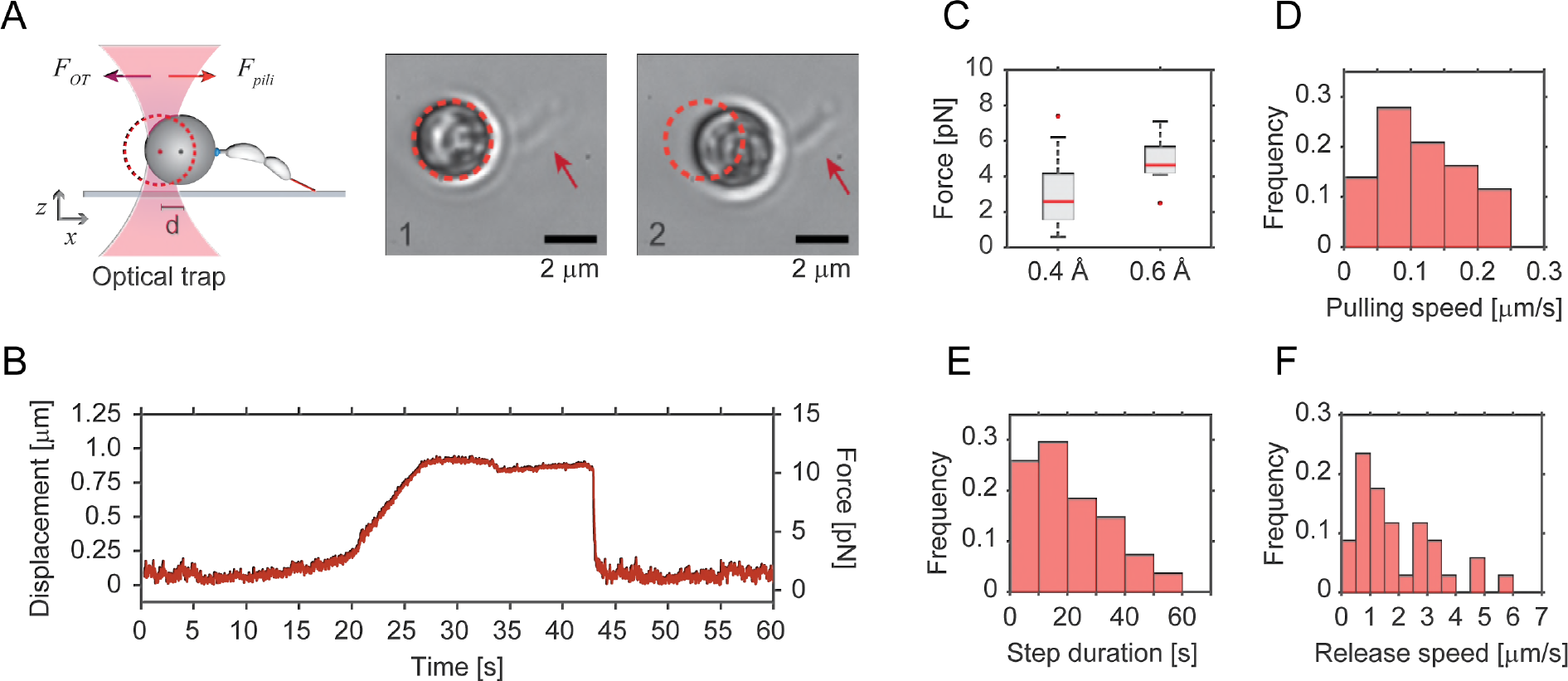
Optical tweezers Determine pili retraction force and speed. **A -** Schematic representation of the experimental setup used for optical trap measurements of pili retraction forces. Beads with late predivisional cells attached via their holdfast were trapped and manoeuvred towards surface to allow pili extending from the opposite pole (red) to attach. Upon pili retraction, bead displacement is measured. Images on the right show a representative example of a bead with one attached predivisional cell (arrow) before (1) and after (2) retraction. Note that the trapped bead is displaced by about 1 μm. **B -** Pili mediated displacement of trapped beads over time. The plot is a representative example of an optical tweezer measurement, showing the displacement and the respective force generated by pili retraction. **C, D -** Force and pulling velocity measurements of pili retraction. Median (red line) and quartiles (boxes) of force generated by pili retraction are shown in (C). Outliers are plotted as red points. The measurements were conducted with the laser power of the optical trap at either 0.4 A (*n* = 100) or at 0.6 A (*n* = 11). The speed of retraction of individual pili is shown in (D) with the laser power set at 0.4 A. **E, F -** The step duration and pili release. Durations of pili retraction (E) were measured as time from the start of bead displacement to the moment of bead release (*n* = 28). (F) The speed with which the bead moves back into the centre oxf the trap at the end of individual step events is indicated (*n* = 34).

**FIGURE 5.**
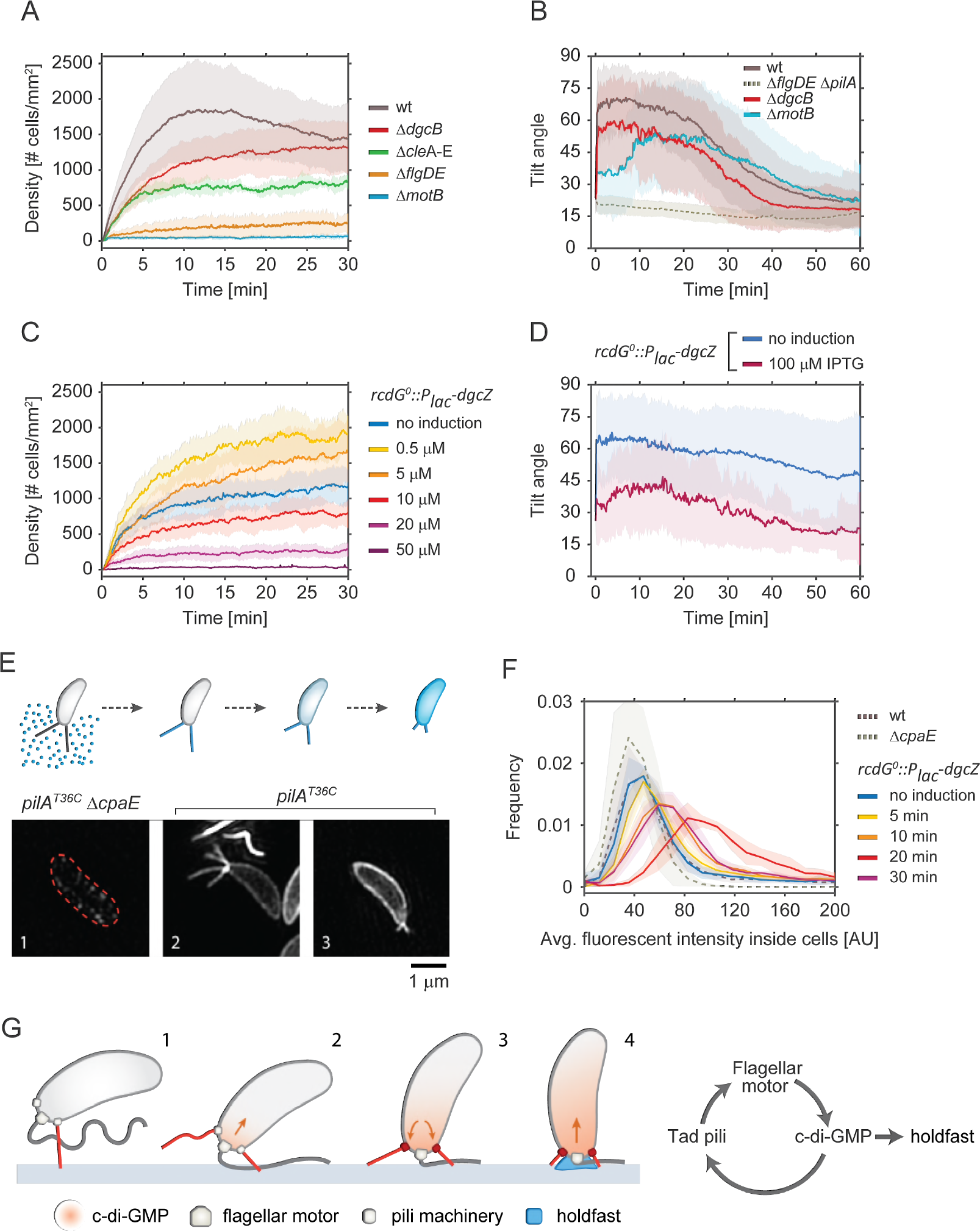
Effect of c-di-GMP on pili activity and surface attachment. **A -** Pili mediated surface attachment in different strains of *C. crescentus* strain. The colonization density was determined over time in a microchannel at constant medium flow rate of 0.75 mm/s. All strains used are defective in holdfast secretion (*hfsA*^−^). Shadow areas represent standard deviations. Replicas: wt = 14, Δ*dgcB* = 14, Δ*flgDE* = 6, Δ*motB* = 10, Δ*cleA-E* = 6. **B -** Pili-mediated standing up of SW cells. The tilt angle ϑ was determined in newborn SW cells of the strains indicated. Time zero corresponds to the moment of SW cell separation from its mother. Shadow areas represent standard deviations. All strains have a functional holdfast. Replicas: wt = 96, Δ*motB* = 54, Δ*dgcB* = 34 and Δ*pilA* = 15. **C -** Pili mediated attachment efficiency as a function of c-di-GMP. Strain NA1000 *hfsA*^−^ *rcdG*^*0*^*::P*_*lac*_-*dgcZ* was grown at increasing IPTG concentration to raise intracellular c-di-GMP and was analysed for surface colonization as outlined in (A). Shadow areas represent standard deviations. Replicas: no induction = 8, 0.5 μM = 6, 10 μM = 6, 20 μM = 4, 50 μM = 6. **D -** Pili-mediated standing up of SW cells as a function of c-di-GMP concentration. The angle ϑ was determined for new-born SW cells of strain *rcdG*^*0*^*::P*_*lac*_-*dgcZ* after cell division. Two distinct *dgcZ* expression levels were tested: no induction (*n* = 44) and 100 μM IPTG (*n* = 41). Shadow areas represent standard deviations. **E -** The upper schematic shows the labelling process for pili. Fluorescent maleimide dye in the medium can label only exposed cysteines, such as those in elongated pili carrying mutated pilin subunits *pilA*^T36C^. When labelled pili are retracted pilin subunits diffuse in the cell membrane and the fluorescent signal correlates with the amount of pilin subunits in the membrane. The lower part of the figure shows representative super resolution images of labelled cells. (1) Strains that carry *pilA*^T36C^, but are unable to assemble pili due to a mutation in the motor machinery (Δ*cpaE*) do not acquire any significant fluorescence. (2) In cells carrying *pilA*^T36C^ and a functional machinery, elongated pili are strongly labelled. (3) Upon retraction disassembled pilin subunits reside in the membrane. **F -** The chart shows the distribution of the average fluorescent signal of SW cell membranes in different strains carrying the mutation *pilA*^T36C^ after labelling with AF488-mal for exact incubation time windows. (Replicas > 3; analysed cells > 6000). Note that because AF488-mal non-specifically binds to holdfast material, all strains used here were devoid of holdfast. **G -** Model of surface attachment and transition from temporary to long term attachment. Within few seconds of landing, cells sense the surface via the flagellar motor and increase their levels of intracellular c-di-GMP. In turn c-di-GMP triggers the secretion of the holdfast and increases the retraction rate of pili.

To avoid potential interference from the flagellum, these measurements were carried out with a strain lacking the external parts of the rotary motor (Δ*flgDE*). A representative example of an optical trap measurement is shown in Figure 4B. Initially, the bead is resting in the center of the optical trap, with some noise due to Brownian motion. Upon pilus attachment to the surface, Brownian motion decreases followed by bead displacement of up to 1 μm from the center before rapidly moving back into its original position (Figure 4B). Bead displacement (*d*) and trap stiffness (*K*_*trap*_) allowed calculating the maximum retraction force (*F* = *K*_*tra*_·*d*). We tested different *K*_*trap*_ and found a maximum retraction force applied by pili during retraction of approximately 8 pN (Figure 4C). This value is close to the approximate drag force of 10 pN that surface bound cells experience in a microchannel with flow speed of 0.75 mm/s (Supplemental Figure 1B). The average speed of pili retraction was 100 nm/s (Figure 4D). In contrast, the speed at which beads backtracked into the trap center was considerably higher (Figures 4E,F), arguing that these events resulted from pili detachment rather than re-elongation of the filaments. From this, we concluded that elongation events are unlikely to interrupt retraction phases, suggesting that pilus retraction is processive disassembling without interruption until all pilin subunits are internalized.

Examining TEM images of piliated cells revealed an average length of pili *L*_*pili*_ = 1.14 ± 0.92 μm (Supplemental Figure 3A). Full retraction of a pilus under flow performs a work equivalent to *W* = 0.5·*K*_*trap*_·*L*_*pil*_^2^ = 4.5·10^−18^ J (or 4.5 10^3^ pN·nm). Assuming 100% efficiency in energy conversion during ATP hydrolysis (80 pN-nm per ATP), we estimated that the pilus machinery could disassemble *L*_*pili*_ / (*W* / *E*_ATP-hydrolysis_) = 20 nm of the filament for each ATP hydrolyzed. A structure-based model for pili Type IV assembly suggested that the machinery uses one ATP subunit to add or subtract one pilin subunit (Chang *et al*, 2017a). In comparison, cryo-EM reconstruction of the gonococcal Type IVb pilus filament revealed a length change of 1.05 nm for each pilin subunit added or removed (Conrad *et al*, 2011). Thus, dividing the length change for ATP hydrolysis at full efficiency (20 nm) by the average length of a pilin subunit (1.05 nm) we estimate the energy conversion efficiency of the *C. crescentus* pilus machinery to be around 5%.

### C-di-GMP regulates Tad pili activity

To better understand the role of pili during surface attachment and their functional interaction with the flagellum, we scored the colonization efficiency of non-motile mutants in microfluidic channels, using the same setup as described above (Figure 2). Both the Δ*motB* and the Δ*flgDE* mutants showed very low levels of colonization densities, suggesting that active swimming is important to efficiently reach the surface in this experimental setup (Figure 5A). Interestingly, non-motile strains also showed higher detachment frequencies and shorter residence times as compared to wild type (Supplemental Figures 4A,B), a behaviour that was most pronounced for the Δ*motB* strain. Since the MotA and MotB stator units of the flagellar motor are involved in surface sensing (Hug *et al*, 2017), this indicated that the flagellar motor and pili may be connected through a regulatory feedback mechanism.

It was recently shown that *C. crescentus* surface sensing leads to a rapid increase of the second messenger c-di-GMP, which in turn mediates biogenesis of the adhesive holdfast (Ellison *et al*, 2017; Hug *et al*, 2017). One of the enzymes implicated in this process is the diguanylate cyclase DgcB (Hug *et al*, 2017). However, it has remained unclear if Tad pili are positioned upstream of c-di-GMP and contribute to the increase of the second messenger during mechanotransduction, or if Tad pili are a target of c-di-GMP control during this process. To clarify this, we monitored pili behaviour in response to changing levels of c-di-GMP. Intriguingly, a Δ*dgcB* mutant showed a significant delay in pili-mediated surface colonization (Figure 5A) and a minor defect in standing up against the medium flow (Figure 5B) as compared to wild type. The fact that in these experiments pili-mediated behaviour is partially affected in the Δ*dgcB* mutant indicated that additional diguanylate cyclase(s) may be involved in this process (Hug *et al*, 2017). This is in line with the observation that new-born SW cells of a Δ*motB* mutant that are unable to sense surface showed a considerably stronger defect in reorienting into an upright position against the media flow (Figure 5B), while a Δ*flgDE* mutant was not affected (Figure 2E). In agreement with the view that Tad pili are able to respond to changes in c-di-GMP, we found that pili-mediated attachment is strongly reduced in a mutant lacking several CheY-like Cle proteins (Figure 5A). Cle proteins bind c-di-GMP to interact with the flagellum and are an integral part of *C. crescentus* surface recognition (Nesper *et al*, 2017).

To analyse the role of c-di-GMP for pili dynamics, we tested a strain harboring an inducible copy of *dgcZ* (*rcdG*^0^*::P*_*lac*_-*dgcZ*). While low level induction of *dgcZ* increased pili-mediated surface attachment, surface colonization gradually decreased upon stronger induction of *dgcZ* (Figure 5C). This effect was the consequence of decreased attachment frequencies (Supplemental Figure 4A). Low level induction of *dgcZ* also decreased detachment rates and increased the cells’ residence time on surface, effects that were reversed at higher induction of *dgcZ*. Similarly, increased induction of *dgcZ* also strongly interfered with the ability of new-born SW cells to stand up (Figure 5D, Supplemental Figure 4C). These observations are in line with an earlier report demonstrating that c-di-GMP is strictly required for *C. crescentus* Tad pili expression and assembly (Abel *et al*, 2013). Transmission electron microscopy revealed that the number of pili per SW cell increased upon surface exposure and that piliation was indeed dependent on the intracellular c-di-GMP concentration. While low level induction of *dgcZ* increased piliation, the number of pili per cell decreased at higher *dgcZ* expression levels (Supplemental Figure 4D). Together, these results indicated that both the flagellar motor and c-di-GMP control Tad pili-mediated behaviour, arguing that c-di-GMP is positioned upstream of pili in the mechanotransduction pathway and is able to regulate pili dynamics.

### C-di-GMP promotes Tad pili retraction

The above results suggested that high levels of c-di-GMP promote Tad pili retraction. To directly visualize Tad pili dynamics, we used a *pilA*^T36C^ to fluorescently label pili subunits with maleimide-based fluorescent dyes. This technique was recently used to visualize Tad pili retraction in real time (Ellison *et al*, 2017). Although we were able to visualize sporadic events of pili retraction, pili filaments are fragile and are prone to breaking during the labelling process. Because only very few cells showed dynamic pili after labelling, it was difficult to obtain statistically solid data sets to compare the behaviour of different mutant strains. To overcome this problem, we adopted a more static fluorescence assay. Because the fluorescent dyes are unable to penetrate the cell envelope, PilA subunits are only labelled if they are assembled into a pilus filament, thereby crossing the outer membrane (Ellison *et al*, 2017) (Figure 5E). Upon pili retraction, disassembled PilA subunits diffuse back into the cytoplasmic membrane thereby increasing the fluorescence signal in this compartment (Figure 5E) (Ellison *et al*, 2017). Thus, to be able to quantify pilus retraction we determined the fluorescence intensities of cell membranes rather than directly monitoring the fragile exterior pili filaments. Importantly, cells expressing wild-type *pilA* or cells expressing *pilA*^T36C^ but lacking the motor subunit of the pilus machinery (Δ*cpaE*) did not accumulate fluorescence. In contrast, cells expressing *pilA*^T36C^ showed a weak but robust fluorescence signal (Figures 5E,F, Supplemental Figure 4E). Similarly, a strain harboring *dgcZ* (*rcdG*^0^*::P*_*lac*_-*dgcZ*) showed weak fluorescence when the expression of the exogenous diguanylate cyclase was not induced (Figure 5F and Supplemental Figure 4E). To monitor Tad pili dynamics at increasing c-di-GMP concentrations, *dgcZ* expression was induced for increasing amounts of time before external pili were labelled by adding the dye. When *dgcZ* expression was induced for 5 minutes, the average fluorescence intensity of cells slightly increased as compared to uninduced cells. Induction of *dgcZ* for 10 and 20 minutes, respectively, gradually increased the amount of fluorescence accumulating in the cytoplasmic membrane in a high proportion of cells (Figure 5F and Supplemental Figure 4E). Thus, as c-di-GMP levels build up, Tad pili activity increases. However, when *dgcZ* was induced for more than 20 minutes before pili were labeled by externally adding the fluorescent dye, fluorescence in the cytoplasmic membrane was reduced again. Importantly, *dgcZ* induction did not affect the overall concentration of pilin subunits (Supplemental Figure 4F). These results support a model where moderate c-di-GMP levels increase pili dynamics while higher c-di-GMP levels lead to the retraction of Tad pili (Figure 5G).

## DISCUSSION

Bacteria have evolved complex mechanisms that enable them to effectively colonize surfaces through a combination of tactile sensing and the exposure of surface adhesins. We have shown earlier that *C. crescentus* SW cells are able to sense surface encounter with their polar flagellar motor and, in response, deploy an adhesive holdfast to remain irreversibly anchored on the surface. Based on the results presented here, we propose that the spatially associated flagellum and Tad pili synergistically optimize the *C. crescentus* surface response. The succession of events leading from temporary to long-term attachment is summarized in the model in Figure 5G. SW cells swimming in close proximity to a surface are able to attach via pre-existing Tad pili (1), thereby creating opportunities for initial surface sensing by the flagellar motor (2). Activation of the motor-coupled diguanylate cyclase DgcB (and possibly other diguanylate cyclases) then generates an increase of c-di-GMP, which boosts the activity of Tad pili. The assembly of additional polar pili increases the cell’s probability to remain attached to and to form tight connections with the surface (3). As c-di-GMP levels increase, processive pili retraction shoves the flagellated pole into close proximity and collision with the surface. In particular, re-orientation of cells into an upright position optimally positions the flagellar motor, strengthening tactile sensing and further increasing c-di-GMP to peak levels required for the allosteric activation of holdfast biogenesis (4) (Hug *et al*, 2017). Our model proposes that the role of pili goes beyond that of passive adhesins promoting temporary attachment and that, instead, Tad pili and the flagellar motor together promote surface adherence in a highly dynamic and coordinated fashion. The model predicts that Tad pili actively guide the efficient surface sensing by the flagellar motor and by that contribute to the critical upshift of c-di-GMP. This model is compatible with the notion that Tad pili themselves may contribute to surface sensing during this process (Ellison *et al*, 2017).

Evidence for our model comes from the direct observation of Tad pili, which increase in numbers when cells are surface exposed or when they experience a moderate increase of c-di-GMP, but decrease in numbers when c-di-GMP reaches peak levels. Moreover, incorporation of fluorescently labeled pilin subunits indicated high Tad pili activity at intermediate levels of c-di-GMP but reduced pili activity when levels of c-di-GMP further increased. Finally, pili-mediated surface attachment and vertical positioning of cells was abrogated when c-di-GMP levels were artificially increased. Based on these observations, we speculate that at moderate levels of c-di-GMP the overall number of polar pili per cell increases through a boost in assembly, slowed disassembly or a combination thereof. In contrast, peak levels of c-di-GMP, which are reached upon sustained surface sensing or during development of the motile SW into a sessile ST cell (Paul *et al*, 2008; Abel *et al*, 2013), signal pili retraction and internalization. Thus, the functional interdependence of the motor, c-di-GMP and pili retraction generates a positive feedback loop that imposes a ratchet-like process that progressively drives cells towards permanent surface attachment (Figure 5G). We had shown earlier that pili are not required *per se* for the motor-mediated surface response and that bacteria grown in very narrow microfluidic chambers where they constantly encounter surface, are able to rapidly attach via their holdfast even if they lack pili (Hug *et al*, 2017). However, under normal conditions bacteria swimming close to a substratum experience surface contact only transiently, a situation that may fashion the need for a highly dynamic and efficient process to stabilize and reinforce this interaction on short time scales. Pili colocalize with the flagellum at one cell pole and are thus optimally positioned to direct the motor towards the surface and to incite collisions that optimize strength and duration of mechanosensing. Tad pili are present (Skerker & Shapiro, 2000) and active (Figure 2) in the predivisional cell immediately before cell division takes place at a time when the flagellum becomes fully operational (Hug *et al*, 2017). In line with the idea that pili reinforce the motor-mediated surface program, their activity in the predivisional cell strictly correlates with the ability of SW offspring to permanently attach next to their mothers in strong flow (Figure 2). Likewise, SW cells that freely rotate around their long axis before being separated from their mothers, presumably because they fail to be dragged towards the surface by Tad pili, rarely manage to remain attached to the surface. In contrast, cells that stopped their rotation generally remained attached after budding off their mothers (Hug *et al*, 2017). These observations directly link pili activity with flagellar obstruction and surface sensing.

The notion that c-di-GMP, depending on its concentration, influences pili activity in distinct ways is supported by observations that c-di-GMP promotes Type IV pili assembly in several bacteria, including *C. crescentus* (Abel *et al*, 2013), *V. cholerae* (Jones *et al*, 2015) and *P. aeruginosa* (Jain *et al*, 2017; Laventie *et al*, 2018). Moreover, increased and decreased levels of c-di-GMP were shown to impact Type IV pili based motility in *Myxococcus xanthus*, arguing that the second messenger is required for but interferes with pili function at increased concentrations (Skotnicka *et al*, 2016). Similarly, polar Tad pili are retracted during the *C. crescentus* SW-to-ST transition coincident with c-di-GMP reaching peak levels (Skerker & Berg, 2001; Abel *et al*, 2013; Paul *et al*, 2008). Mechanistically, this complex regulation could result from the antagonistic activities of two effector proteins that bind c-di-GMP with different affinities to promote the assembly and disassembly of pili filaments, respectively. Similar mechanisms were described in *P. aeruginosa* and *X. campestris*, where two c-di-GMP binding proteins, FimX and FimW, localize to the cell poles to modulate pili formation. While the mode of action of FimW is still unknown, FimX facilitates pili elongation during twitching at the leading cell pole by interacting with the assembly motor ATPase PilB (Guzzo *et al*, 2009; Jain *et al*, 2017). Direct interaction of c-di-GMP with the pili machinery was also shown in *V. cholerae*, where c-di-GMP binding by the motor assembly protein MshE promotes polymerization of pili in a dose dependent manner (Jones *et al*, 2015). How exactly c-di-GMP influences pili assembly and retraction in *C. crescentus* remains to be shown.

Our model for *C. crescentus* surface attachment proposes tight functional cooperation of the flagellar motor and Tad pili. Evidence for this stems from the observation that pili activity was reduced in mutants lacking the MotB stator unit or lacking all five Cle proteins (CleA-E) (Figure 5A,B). *C. crescentus* Cle (CheY-like) proteins were recently shown to bind c-di-GMP and, in response, interact with the flagellar motor to impede flagellar activity and promote surface adaption (Nesper *et al*, 2017). It was proposed that one or several of these proteins is part of a positive feed-back loop that reinforces the motor response during surface sensing thereby facilitating a rapid upshift of c-di-GMP and production of holdfast material. The reduced pili-mediated attachment observed for the Δ*cleA-E* strain could thus be due to an insufficient raise in c-di-GMP or, alternatively, may implicate one of the Cle proteins in the regulation of pili directly, providing a mechanistic basis for the coordination of both organelles. Further studies are needed to clarify the role of Cle components in this process.

Experiments in flow devices demonstrated that *C. crescentus* Tad pili are highly dynamic and are able to promote twitching- or walking-like movements of *C. crescentus* SW cells (Figure 3). For these experiments we used a mutant lacking the HfsK N-acetyltransferase, an enzyme that was proposed to chemically modify holdfast material. A Δ*hfsK* mutant forms malleable holdfast structures that can cling cells to surfaces but lack the adhesive strength required to hold cells in place under flow or when pulled by other forces like pili retraction. It was recently proposed that HfsK acylates the EPS component of the holdfast and that this modification is necessary for proper holdfast cohesion and anchoring (Sprecher *et al*, 2017). The observation that HfsK activity itself is regulated by c-di-GMP indicated that under certain conditions holdfast material may be formed, but remain in a non-acylated form. If so, limited cohesive strength of the holdfast could attach *C. crescentus* cells to surfaces without restricting their pili-mediated motility. This would allow this aquatic organism to explore liquid exposed surfaces with their pili motors similar to the well-known twitching and walking motility of soil bacteria.

## SUPPLEMENTAL MATERIAL

**Supplemental Figure 1.**
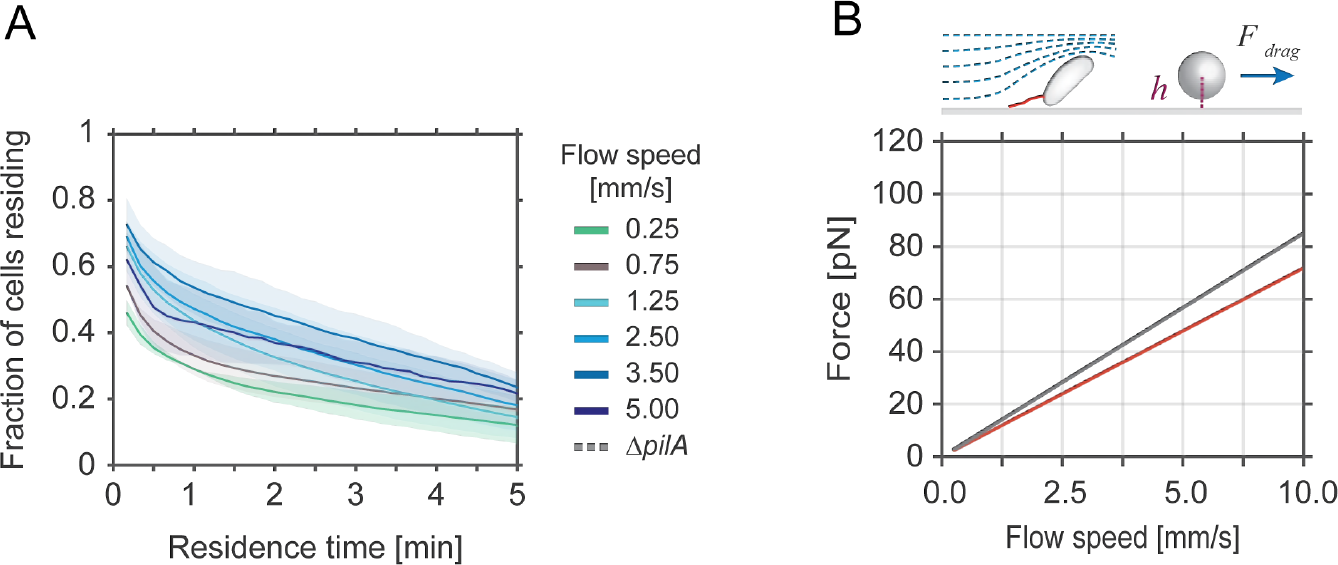
**A -** Residence time of cells on a surface during pili-mediated attachment. Each curve indicates the cumulative fraction of cells residing on surface for a period equal or greater than the indicated time. The *C. crescentus hfsA*^−^ mutant used in these experiments was tested at different flow velocities. Opaque areas represent standard deviations (*n*>3). **B -** Theoretical drag force experienced by a typical SW (red) and ST cell (grey) attached to a surface at different flow rates. The drag force is calculated using the equation shown below and the bacterial shape was simplified as a sphere near a surface with a volume equivalent to the average cell. The drag was calculated assuming a distance from the surface (*h*) of 0.25 μm.

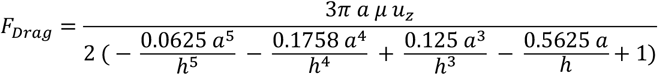 *a* = radius of the sphere with equivalent volume of a cell, being ≈ 0.62 μm^3^ for SW cells and ≈ 0.71 μm^3^ for ST cells, *h* = height from the surface, *μ* = viscosity, *u*_*z*_ = flow velocity at height *h* from the surface. Equation 7-4.37 (Happel & Brenner, 1981).

**Supplemental Figure 2.**
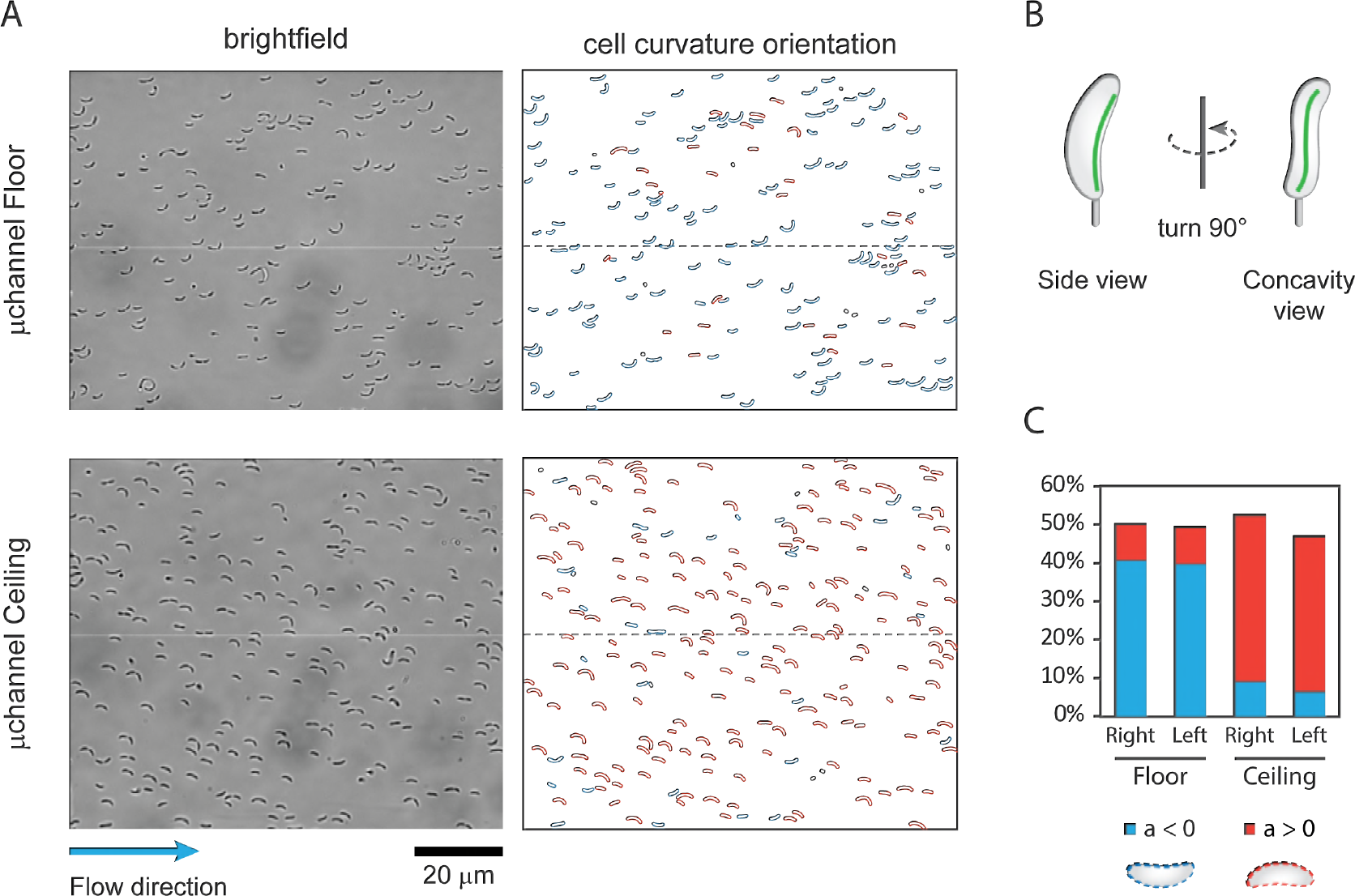
**A -** Brightfield images (left) showing *C. crescentus* cells attached to the floor and ceiling of a microfluidic channel. Images were analysed with a MatLab-based script to determine the cells’ concavity orientation (right). Cells with the concavity oriented to the right (red) or to the left (blue) with respect to the direction of the flow are highlighted. **B -** Schematic drawing of the proposed cell shape of crescentoid *C. crescentus* with an exaggerated left-handed twist. **C -** Quantitative analysis of the concavity orientation (n = 3) with cells scored in an area of ≥ 0.3 mm^2^ both on the ceiling and floor surfaces of a device. Concavity distribution is identical in the right half (Right) and the left half of the channel (Left) excluding flow related effects.

**Supplemental Figure 3.**
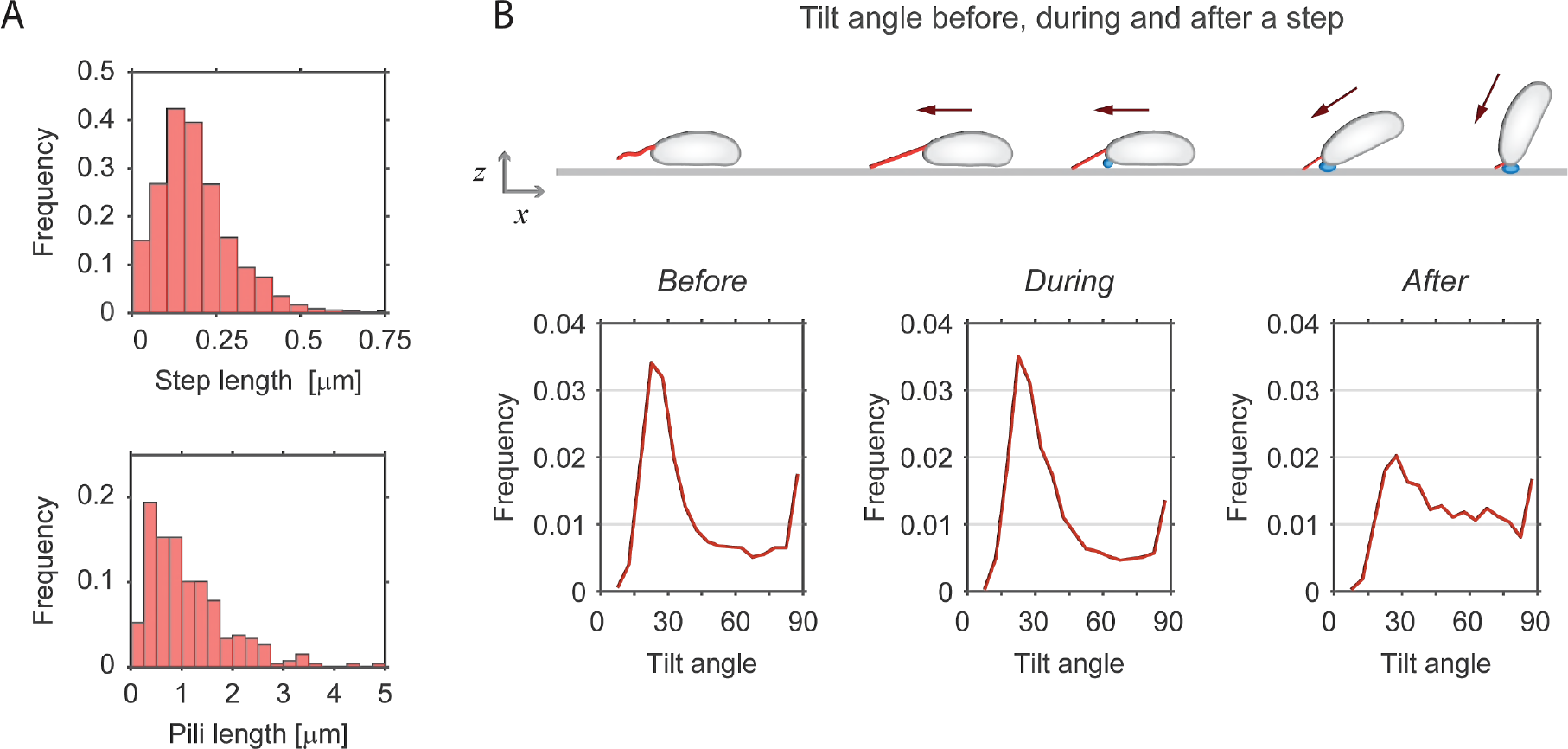
**A -** Step length distribution of *C. crescentus* Δ*hfsK* mutant cells during surface motility (*n* = 56) (upper panel) and distribution of pili length in wild type cells as observed by TEM (*n* = 271) (lower panel). **B -** Schematic of a *C. crescentus* SW cell moving against the medium flow and standing upright at the end of each dislocation step. Pili (red), holdfast (blue) and cell movement (red arrow) are indicated. The charts below the graph show the distributions of tilt angle values five seconds before (left), during (middle) and five seconds after (right) an upstream step event. Before and during a step event cells are more likely to lie flat on the surface, while standing up upon completion of an upstream movement.

**Supplemental Figure 4.**
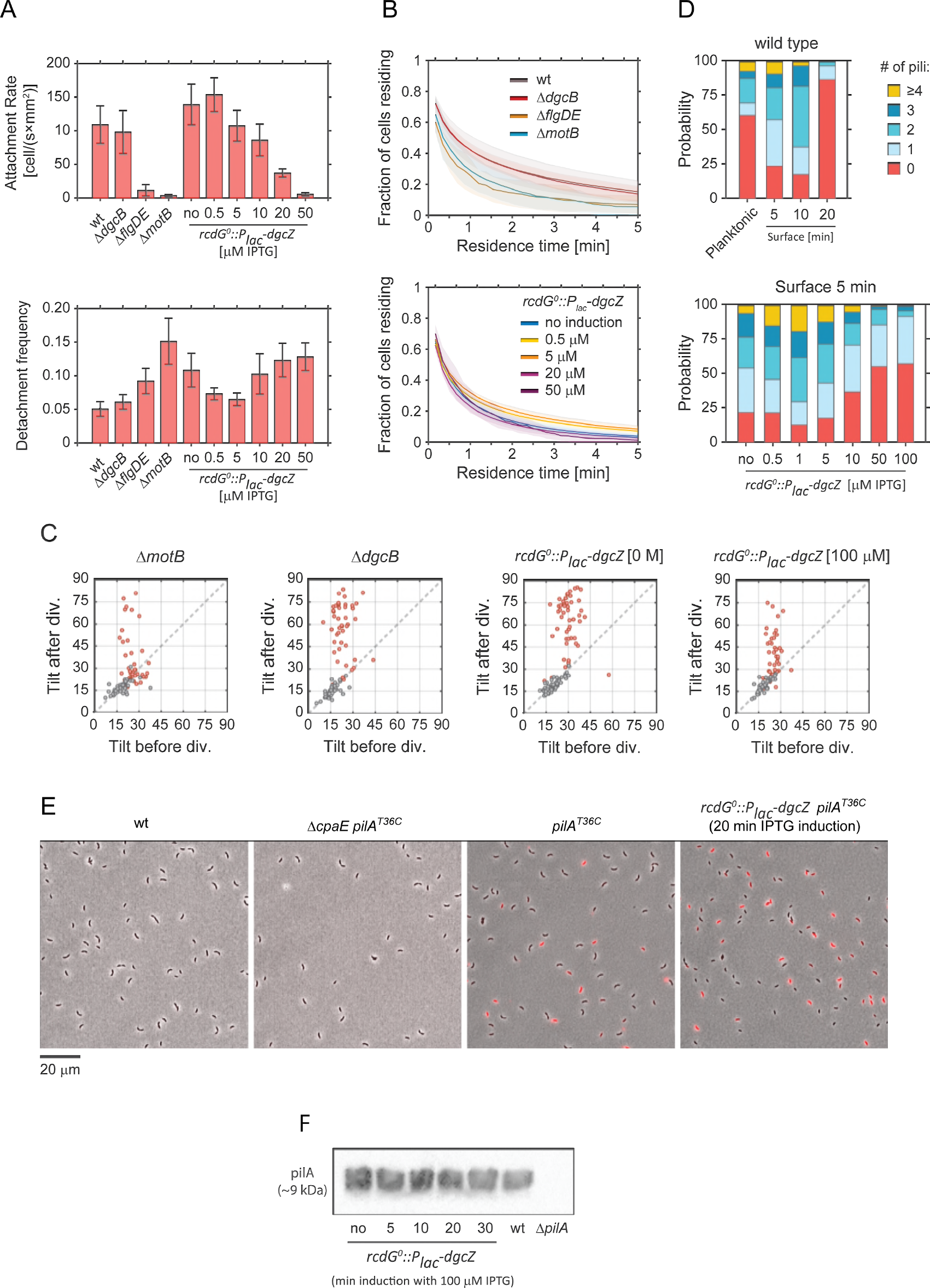
**A -** Surface attachment of SW cells of different wild-type and mutant strains in microfluidic devices. Average numbers of newly attached cells per mm^2^ per second are shown in the upper panel. The lower panel shows desorption frequencies of the same strains, calculated as the ratio of cells leaving the surface over the total number of cells attached between two time points (5 s). Values were obtained from the attachment assays shown in Figure 5A and C during the time window between minutes 10 and 25. Error bars indicate standard deviations. **B -** Residence time of cells on surface during pili-mediated attachment. Each curve indicates the cumulative fraction of cells residing on surface for a period equal or greater than the indicated time. Opaque areas represent standard deviations. All strains are unable to secrete holdfast (*hfsA*^−^). Replicas: upper chart > 5; lower chart > 4. **C -** Scatter plots with the average angle Ñ of SW (red) and ST cells (grey) recorded 5 minutes before and 5 minutes after cell separation. Replicas: Δ*motB* = 41; Δ*dgcB* = 45, *rcdG*^0^::*P*_*lac*_-*dgcZ* (0 M) = 50; *rcdG*^0^::*P*_*lac*_-*dgcZ* (1 μM) = 46. **D -** Number of pili observed at the pole of individual *C. crescentus* wild-type cells imaged by TEM. In the upper chart, wild-type cells were either fixed before (planktonic) or after being spotted on EM grids for 5, 10 and 20 minutes (surface) to allow them to make surface contact. The fraction of cells with specific numbers of pili are indicated. The lower chart shows pili numbers in the strain *rcdG*^0^::*P*_*lac*_-*dgcZ* at different levels of IPTG induction. In this case, cells were fixed 5 minutes after making surface contact. **E -** Representative images of different *C. crescentus* strains after pili labelling. Strains engineered to express the *pilA*^*T36C*^ allele were specifically labelled with the fluorescent dye AF-647-mal. Strains expressing a wild-type pilA allele or defective in pili assembly (Δ*cpaE*) were used as controls. Images were acquired by fluorescence microscopy. Bright field (grey-scale) and fluorescence images are overlaid (red). The fluorescent channel of the different images was set to the same parameters. All strains used were unable to secrete holdfast (*hfsA*^−^). **F -** Immunoblot analysis of *C. crescentus* wild type and mutant strains using an antibody against the major pilin subunit PilA. The strain *rcdG*^0^::*P*_*lac*_-*dgcZ* was tested without IPTG induction or in the presence of 100 μM IPTG for different time windows. *C. crescentus* wild type (wt) and Δ*pilA* mutant samples are used as controls.

**Supplemental Table S1.** Bacterial strains and Plasmids used in this study.

| ID | Name | Description | Reference |
| --- | --- | --- | --- |
|  |  | *C. crescentus* NA1000 *hfsA*<sup>-</sup> |  |
| UJ4475 | *hfsA*<sup>-</sup> | NA1000 is a wild type strain laboratory strain derived from CB15. It lacks holdfast expression due to a mutation in *hfsA*. | (Evinger & Agabian, 1977) |
| UJ5511 | *hfsA*<sup>+</sup> | Wild type *hfsA* gene was restored for this strain. | this work |
| UJ8177 | $\Delta flgDE$ (with holdfast) | Clean deletion of flagellar hook *flgD* and *flgE* genes, Filament-less flagellum. Non-motile. | this work |
| UJ3299 | $\Delta flgDE$ | Clean deletion of flagellar hook *DflgD* and *flgE* genes. Filament-less flagellum. Non-motile. | this work |
| UJ8178 | $\Delta flgDE \Delta pilA$ (with holdfast) | Clean deletion of flagellar hook *flgD* and *flgE* genes, and *pilA* gene. Filament-less flagellum. Non-motile. Pili negative strain. | this work |
| UJ7040 | $\Delta pilA$ | Clean deletion of the *pilA* gene. Pili negative strain. | (Nesper *et al*, 2017) |
| UJ10591 | $\Delta creS$ (with holdfast) | Clean deletion of the *creS* gene. Strain with rod-like cell shape instead of the typical curvature. | this work |
| UJ8179 | $\Delta motB$ (with holdfast) | Encoding *motB_D33N* variant at the native locus. Stator without proton conductance. Non-motile flagellum. | (Hug *et al*, 2017) |
| UJ10250 | $\Delta motB$ | Encoding *motB_D33N* variant at the native locus. Stator without proton conductance. Non-motile flagellum. | this work |
| UJ9064 | $\Delta dgcB$ (with holdfast) | Encoding *dgcB_E216Q* variant at the native locus. The cyclase is catalytically inactive. | (Hug *et al*, 2017) |
| UJ7154 | $\Delta dgcB$ | Encoding *dgcB_E216Q* variant at the native locus. The cyclase is catalytically inactive. | this work |
| UJ7853 | $\Delta cleA-E$ | Clear in frame deletion of *cc0440*, *cc1364*, *cc2249*, *cc3100* and *cc3155*. | (Nesper *et al*, 2017) |
| SoA1587 | *hfsA*<sup>-</sup> $\Delta dgcA \Delta dgcB \Delta pleD \Delta pdeA \Delta cc0091 \Delta cc0655 \Delta cc0740 \Delta cc0857 \Delta cc0896 \Delta cc1086 \Delta cc3094 \Delta cc3148 P_{lac}-dgcZ$ | rcdG<sup>0</sup> strain. Clean deletions of all endogenous GGDEF and EAL-domain encoding genes in NA1000. Chromosomal integration of *E. coli dgcZ-3xflag* gene under the control of the Plac promoter. | (Abel *et al*, 2013) |
| UJ7771 | *hfsA*<sup>-</sup> $\Delta dgcA \Delta dgcB \Delta pleD \Delta pdeA \Delta cc0091 \Delta cc0655 \Delta cc0740 \Delta cc0857 \Delta cc0896 \Delta cc1086 \Delta cc3094 \Delta cc3148 P_{lac}-dgcZ$ (with holdfast) | SoA1587 strain with restored *hfsA* gene. | (Hug *et al*, 2017) |
| UJ7604 | $\Delta dgcA \Delta dgcB \Delta pleD \Delta pdeA \Delta cc0091 \Delta cc0655 \Delta cc0740 \Delta cc0857 \Delta cc0896 \Delta cc1086 \Delta cc3094 \Delta cc3148 P_{lac}-dgcZ \Delta hfsK$ (with holdfast) | UJ7771 with clean deletion of gene *hfsK* (*cc3689*), which lowers cohesion of the holdfast. | this work |
| YB8288 | *pilA*<sup>T36C</sup> | Encoding *pilA_T36C* variant at the native locus. The protein can be labelled with maleimide-reactive dye. | (Ellison *et al*, 2017) |
| UJ10592 | *pilA*<sup>T36C</sup> Δ*cpaE* | YB8288 with clean deletion of *cpaE* gene. Pili negative strain. | this work |
| UJ10321 | *pilA*<sup>T36C</sup> *rcdG*<sup>0</sup>::*P<sub>lac</sub>-dgcZ* | SoA1587 strain encoding *pilA_T36C* variant at the native locus. | this work |

| ID | Plasmids | Description | Reference |
| --- | --- | --- | --- |
| pMS100 | pNPTS139 *PilA*<sup>T36C</sup> | Used to introduce *pilA_T36C* variant at the native locus. | this work |
| pSA223 | pNPTS138 - *xylR<sub>up</sub>-T7-term- P<sub>lac</sub>-dgcZ-3xflag* | Used for introducing exogenous cyclase *dgcZ* into the genome. | (Abel *et al*, 2013) |

## METHODS

### Bacterial strains and growth conditions

The wild type (*wt*) strain of reference was either NA1000 *hfsA*^−^, a lab adapted strain with a point mutation in gene *hfsA*, which makes it unable to secrete the holdfast, or NA1000 *hfsA*^+^, where the functional gene was reintroduced. The strain *rcdG*^0^ is a c-di-GMP-free strain where all major endogenous GGDEF and EAL-domain encoding genes were deleted. Introducing in *rcdG*^0^ the exogenous diguanylate cyclase *dgcZ* under the control of the IPTG-inducible *lac* promoter (*P*_*lac*_-*dgcZ*), allows to tune the expression of *dgcZ* and in turn to control the intracellular levels of c-di-GMP, as previously reported (Abel *et al*, 2013). A full list of strains used in this study is listed in Supplemental Table 1.

### Fabrication of microfluidic devices

Masters were fabricated via standard photolithography protocols (Deshpande & Pfohl, 2012). PDMS (polymethyldisiloxane, Sylgards 184, Dow Corning) devices were created via replica molding, and then aged via heat treatment on a hotplate at 150°C for 30 min (Göllner *et al*, 2016). Holes were drilled at inlet(s) and outlet(s), devices were treated in oxygen plasma and covalently bound onto borosilicate cover glass slides (round, 50 mm diameter, thickness No. 1, VWR). All microflow experiments were conducted in single microchannels with dimensions of either of μm in width and 100 μm in height or a width of 50 μm and a width of 200 μm.

### Microfluidics and microscopy setup

For attachment assays overnight cultures (generally *hfsA*^−^) were diluted 1:50 and grown at 30°C in PYE medium under agitation. For *rcdG*^0^::*P*_*lac*_-*dgcZ* we added the desired IPTG concentration during the dilution step. Once the culture reached an OD_660_ of 0.15, it was loaded in a 1 ml plastic syringe (Soft-ject, Henke-Sass, Wolf). The syringe was then plugged to a needle (23G, 0.6 × 30 mm, BraunMelsungen AG) which in turn was connected to one end of a PTFE microtube (0.56 × 1.07 mm, Fisher Scientific). The tubing was filled with the culture and the terminal end of the tubing connected to the inlet of a microfluidic device (channel cross section: 50 μm width by 200 μm height). The syringe was mounted on a syringe pump (neMESYS low pressure module V2, 14:1 gear; Cetoni GMBH). With the microfluidic setup ready, the device was places on the stage of an inverted microscope (IX81, Olympus GMBH). The syringe pump was initially set to create a strong flow of 25 mm/s for 1 min to ensure that the microchannel surface was devoid of cells. Flow was then set to the desired target velocity for a duration of the experiment, 30-45 minutes, at room temperature. Time-lapse images were recorded at 0.1-0.16 fps, using a 40x oil immersion objective (UPlanFLN 40x Oil, Olympus). We used a new batch of culture at OD_660_ 0.15 for each experiment.

For observations of single cell division events, a culture of the desired strain (generally *hfsA*^+^) was loaded in the microfluidic channel of a device (channel cross section: 25 μm width by 100 μm height). Cells were left to colonize the surface for a short period resulting in an average distance between cells of about 20-30 μm. This low density ensured that single cell division events were not disturbed or influenced by neighboring cells. A 1 ml plastic syringe loaded with fresh PYE medium (plus the desired IPTG concentration for *rcdG*^0^::*P*_*lac*_-*dgcZ*) was plugged to a needle and connected to one end of the a PTFE microtube. The syringe was then mounted on a syringe pump and the terminal end of the tubing plugged into the device’s inlet. A steady flow of 1 mm/s was then set for the duration of the entire experiment. Cells were left to growth for 1-2 h before recording started with image sequences at 1 or 5 fps, using a 100x oil immersion objective (CFI Plan Apo λDM100x Oil, Nikon). The experiments were conducted at room temperature, for no more than 10 h. This procedure ensured steady growth conditions and no overgrowth/clogging in the inlet.

### Optical Tweezers setup and force measurements

The optical tweezers experiments were performed on a custom built bright field microscope, complemented with a laser diode setup (LD830-MA1W, λ = 830 nm, Thorlabs). A water immersion, high aperture objective (UPlanSApo 60x water, Olympus) was used to focus the laser beam, trap the beads with attached bacteria, and image the fluctuations of the bead in the trap and the attached bacteria. The experiments were carried out in position clamp mode at a constant laser power. The images of the beads and the attached cells were recorded with 50 Hz and 75 Hz using a fast camera (Phantom Miro EX4, Vision Research Inc.). From the movies we measured the change in position of the bead during the experiment and we observed the active cell.

Calibration of the optical tweezer was carried out via fluctuation calibration. For each laser power used, an image sequence of a bead in the trap was recorded at 1000 Hz. From the fluctuation of the bead in the trap the variance σ was determined. The trap stiffness *K*_*trap*_ was calculated as: *K*_*trap*_ = *k*_*B*_*T*/σ^2^, with *k*_*B*_ the Boltzmann constant and *T* the room temperature.

An exponential culture of NA1000 *hfsA*^+^ Δ*flgDE* at OD_660_ of 0.15 was mixed with polystyrene beads (Fluoresbrite YG Carboxylate Microspheres 3.0 μm, Polysciences) to a final concentration of 1.7·10^8^ beads per mL cell suspension. The mix was incubated for 2 min to allow cells to attach to the beads. Then a 1:1 dilution with fresh PYE was injected into the device and experiments were conducted in no-flow conditions. Devices used for optical tweezer measurements had chambers attached to a main channel (Deshpande & Pfohl, 2012, 2015) connected by an opening of less than 10 μm. Single bead-carrying predivisional cells were chosen and placed inside the chambers where the optical tweezer measurements could be performed undisturbed from other cells.

### Pili staining and fluorescence

The labelling of pili was done following the protocol published in a recent study of Ellison C.K. et al (Ellison *et al*, 2017). Briefly, an overnight culture was diluted 1:50 or 1:100 and grown to an OD_660_ of 0.10, under agitation. 1 ml of culture was transferred into an Eppendorf vial and 25 μg/ml of maleimide-reactive dye AF647-mal (Sigma-Aldrich) were added and gently mixed by inverting. The sample was incubated for exactly 5 min and was then centrifuged (4000 g for 1 min), washed with 1 ml fresh PYE and centrifuged again. The final pellet was resuspended in 50 μl fresh PYE. Finally, 2 μl were spotted on a 1% agarose pad and imaging at the microscope was conducted immediately. For *rcdG*^0^::*P*_*lac*_-*dgcZ*, we added 100 μM IPTG for an exact time window before the washing step: the total duration of IPTG induction included the 5 minutes maleimide-labelling step. For induction times longer than 5 min, the culture was kept in incubation until the labelling step.

The time elapsed between washing step and beginning of imaging was minimized, on average taking 4-5 min. Each sample was imaged for no longer than 7-8 minutes after the labelling step. Imaging was done at an inverted microscope (Eclipse Ti2, Nikon Instruments Europe B.V.) using a 100x oil immersion objective (CFI Plan Apo λDM100x Oil, Nikon). We imaged 100-150 random positions for each pad and for each position we acquired a phase contrast (exposure 50 ms) and a fluorescent image (Ex laser 640 nm at 25% intensity for 150 ms, Em mCherry 592-667)

To analyse the fluorescent signal inside cells’ bodies, we used the Matlab-based program microbeTracker for cell detection (Sliusarenko *et al*, 2011) and an in-house built Matlab-based program to analyse the fluorescent channel. We selected SW cells by using as criteria a body length <2.5 μm.

### 3D-SIM super-resolution microscopy

Three-dimensional structured illumination microscopy (3D-SIM) was performed on a DeltaVision OMX-Blaze V4 system (GE Healthcare). Images were acquired using a Plan Apo N 60x, 1.42 NA oil immersion objective lens (Olympus) and 4 liquid-cooled sCMOS cameras (pco.edge 5.5, full frame 2560 x 2160; PCO). Exciting light was directed through a movable optical grating to generate a fine-striped interference pattern on the sample plane. The pattern was shifted laterally through five phases and three angular rotations of 60° for each z-section. The 488 nm laser lines were used during acquisition and the optical z-sections were separated by 0.125 μm. Laser power was attenuated to 10% and exposure times were typically 80 ms. The laser power was adjusted to achieve optimal intensities between 2,000 and 3,000 counts in a raw image of 15-bit dynamic range at the lowest laser power possible to minimize photobleaching. Raw 3D-SIM images were processed and reconstructed using the DeltaVision OMX SoftWoRx software package (v6.1.3, GE Healthcare).

### Image analysis: cell orientation and tracking

Cell outline detection was carried out using the Matlab-based program microbeTracker (Sliusarenko *et al*, 2011). Further tracking, analysis and statistics were carried out with in-house developed Matlab scripts. Cell perimeters were fitted to an ellipse and the eccentricity of the ellipses (*ε*), which is the ratio between the major and minor axes, were monitored. We used *ε* to calculate the inclination of cells in respect of the surface (angle ϑ). A *ε* value of 1 was set to correspond to the tilt angle *θ* of 90° and a *ε* value of 0.1 to correspond with a tilt of 0°. Eccentricity values below 0.1 could be excluded as they represent very elongated ellipses, with a length-to-width ratio that does not occur for *C. crescentus* SW cells.

To track the trajectories of cells we choose to follow the position of the holdfast. This position lies in a point between a vertex and a focus of the ellipse which fits the 2D projection of the cell. The position was dependent on *ε*, such that as the cell is upright, it coincides with the focus, and as the cell lies flat, it is close to the vertex (see Figure 2C). Step events (see Figure 3B) were identified as sequences of fast movement against the flow, with speeds of ≥0.1 μm/s and sustained for ≥1 second.

### Electron microscopy

To examine surface induced cells, 5 μl sample were spotted on a 400-mesh copper grid covered with a parlodion and carbon film for 5 to 20 min. Then the grid was gently washed using a micropipette and cells were fixed with 0.1% glutaraldehyde. Alternatively, to examine planktonic samples, glutaraldehyde was added to a bacterial liquid culture to a final concentration of 0.1%. Then, 5 μl were spotted on the grid. The samples were washed with water 4 times water and negatively-stained twice with 0.5% uranyl acetate. Images taken at the TEM (Morgagni 268D FEI (80kV) and a FEI T12 (120kV) - TVIPS F416) were visually inspected and SW cells identified by the absence of a stalk and cell length <2.5 μm.

### Immunoblot analysis

Proteins were separated by electrophoresis on 20% SDS-polyacrylamide gels and transferred onto nitrocellulose blotting membrane 0.2 μm (Amersham Protran 0.2 μm NC, GE Healthcare). PageRuler Prestained Protein Ladder (ThermoFisher) was used to mark protein sizes. The primary antibody for detection was polyclonal rabbit anti-PilA (Fumeaux *et al*, 2014), diluted of 1:4000. The secondary antibody was swine anti-rabbit coupled to horseradish peroxidase (Dako), used in dilution of 1:10000. Antibody-treated blots were incubated with LumiGLO (KPL) for exposure of super RX-N films (Fujifilm).

## ACKNOWLEDGEMENTS

We acknowledge Ursula Sauder and Carola Alampi of the C-CINA of Imaging Core University of Basel for technical assistance with the TEM microscopy; the Imaging Core facility (IMCF, University of Basel) and in particular Alexia Ferrand for the technical assistance provided on the OMX microscope; Fabienne Hamburger for plasmid construction and Benoît-Joseph Laventie for fruitful discussions on the manuscript. We thank Yves Brun and Courtney K. Ellison for providing plasmids and strains. We gratefully acknowledge funding by the Swiss Nanoscience Institute in Basel, Switzerland (SNI PhD graduate school, Project P1302) and by the Swiss National Science Foundation (grant 310030B_147090 to U.J.). The authors declare that they have no conflict of interest.

